# Plasma endocannabinoids in cocaine dependence and their interaction with cocaine craving and metabotropic glutamate receptor 5 density in the human brain

**DOI:** 10.1101/2023.04.18.537293

**Authors:** Sara L. Kroll, Lea M. Hulka, Ann-Kathrin Kexel, Matthias Vonmoos, Katrin H. Preller, Valerie Treyer, Simon M. Ametamey, Markus R. Baumgartner, Carola Boost, Franziska Pahlisch, Cathrin Rohleder, F. Markus Leweke, Boris B. Quednow

## Abstract

Animal models indicate that the endocannabinoid system (ECS) plays a modulatory role in stress and reward processing, both crucially impaired in addictive disorders. Preclinical findings showed endocannabinoid-modulated synaptic plasticity in reward brain networks linked to the metabotropic-glutamate-5 receptor (mGluR5), contributing to drug-reinforcing effects and drug-seeking behavior. Although animal models postulate a link between ECS and cocaine addiction, human translational studies are lacking. Here, we tested previous preclinical findings by investigating plasma endocannabinoids (eCBs) anandamide (AEA), 2-arachidonoylglycerol (2-AG), and the related N-acylethanolamines (NAEs) palmitoylethanolamide (PEA) and oleoylethanolamide (OEA), including their interaction with cerebral mGluR5, in chronic cocaine users (CU). We compared basal plasma concentrations between chronic CU (N=103; 69 recreational CU and 34 dependent CU) and stimulant-naïve healthy controls (N=92). Follow-up basal eCB/NAE plasma levels after 12 months were used for reliability and stability check (CU: N=33; controls: N=43). In an additional analysis using ^11^C-ABP688 positron emission tomography (PET) in a male subsample (CU: N=18; controls: N=16), we investigated the relationships between eCBs/NAEs and mGluR5 density in the brain. We found higher 2-AG plasma levels in dependent CU compared to controls and recreational CU. 2-AG levels were stable over time across all groups. In the PET-subsample, a positive association between 2-AG and mGluR5 brain density only in CU was found. Our results corroborate animal findings suggesting an alteration of the ECS in cocaine dependence and an association between peripheral 2-AG levels and cerebral mGluR5 in humans. Therefore, the ECS might be a promising pharmaco-therapeutic target for novel treatments of cocaine dependence.

## Introduction

Impaired reward behavior and inadequate stress response are key features of the development and maintenance of drug addiction. Accordingly, long-term maladaptive functional and structural synaptic plasticity has been reported in brain networks related to reward behavior and stress response [1,2]. Cocaine acts as a non-selective monoamine transporter inhibitor, entailing increased release of dopamine (DA), serotonin, and noradrenaline in the synaptic cleft in animals and humans[3-6]. In particular, the inhibitory effects on the presynaptic DA transporter resulting in elevated DA levels within the brain regions of the mesolimbic reward network (including projections from the ventral tegmental area [VTA] to the nucleus accumbens [NAc]) have been linked to the highly addictive properties of cocaine [7,8]. Animal models have shown that acute cocaine-induced activation of DA neurons in the NAc is specifically associated with the drug-reinforcing effects enhancing motivated and drug-seeking behavior [9-11]. Moreover, preclinical evidence suggests that repeated cocaine use entails drug-induced neuroplastic adaptations in mesolimbic DA neurons, causing drug-cue related hyper-sensitivity and subsequently contributing to cocaine use maintenance and drug relapse even after a prolonged time of abstinence [12-14]. Human neuroimaging and pharmacological studies corroborate animal findings reporting alterations of the mesolimbic DA network in chronic cocaine users (CU) contributing to the drug’s reinforcing and addictive effects [15-24].

In recent years, accumulating evidence mainly from preclinical studies indicates that the endocannabinoid system (ECS) plays a major role in reward processing. Specifically, long-term depression (LTD) causing synaptic plasticity in the mesolimbic DA network has been linked to the ECS underpinning drug-seeking and addictive behavior [25-28]. Animal models consistently showed endocannabinoid-mediated LTD in the VTA and NAc caused by the two main endocannabinoids, anandamide (AEA) and 2-arachidonoylglycerol (2-AG), activating the cannabinoid type-1 (CB_1_) receptor in the mesolimbic reward network. AEA and 2-AG are synthesized postsynaptically on-demand by the primary enzymes N-acylphosphatidylethanolamine specific phospholipase D (NAPE-PLD) and diacylglycerol lipase (DAGL), respectively, and act as retrograde neurotransmitters by activating presynaptic CB_1_ receptor entailing inhibition of presynaptic neurotransmitter release [29-33]. CB_1_ receptors are highly expressed throughout the mesolimbic DA network localized on terminals of glutamatergic and GABAergic neurons but not on DA neurons itself [34-36].

Stimulation of 2-AG-mediated LTD at GABAergic interneurons in the VTA resulting in disinhibition of DA neurons has been shown in animal models entailing increased DA release and projections to the NAc [37-39]. Preclinical findings showed that acute and repeated cocaine administration facilitates endocannabinoid-mediated LTD at GABAergic interneurons and abolishes endocannabinoid-mediated LTD at glutamatergic neurons, both resulting in increased DA release in the VTA and its projection to the NAc, contributing to the drug reinforcing effects [40-42]. Moreover, cocaine-induced activation of the endocannabinoid-LTD mechanism in the VTA has been recently linked to motivational behavior in rats, while the CB_1_ receptor inverse agonist rimonabant was able to block reward-seeking behavior [43]. In the NAc, a single *in vivo* cocaine administration in mice has been shown to abolish endocannabinoid-mediated LTD at terminals of glutamatergic neurons resulting in increased glutamate release and subsequently in increased excitation of DA cells [44]. Synaptic plasticity induced by endocannabinoid-mediated LTD has been further related to activation of postsynaptic metabotropic glutamate receptor 5 (mGluR5) in various brain regions such as the striatum, hippocampus, and amygdala in rodents [45-47]. Animal models showed that activating mGluR5 triggers synthesis and release of 2-AG through the DAGL pathway into the synaptic cleft [48]. Notably, a single administration of cocaine in mice has been shown to alter the functional interaction of endocannabinoid-mediated LTD and mGluR5 [44]. Recently, activation of mGluR5-mediated 2-AG synthesis and release at D1-medium spiny neurons (D1-MSN) in the NAc amplifying LTD at glutamatergic neurons has been linked to reward-seeking behavior due to inhibition of glutamate release [25,49]. Accordingly, genetic deletion of mGluR5 in D1-MSN in mice models abolished 2-AG-mediated LTD, preventing expression of drug and natural reward as well as brain-stimulation-seeking behavior. This mechanism was restored by pharmacological enhancement of 2-AG signaling in the NAc in mGluR5-ablation mice. This leads to the assumption that not exclusively mGluR5 but its specific interaction with 2-AG plays an essential role in reward-and drug-seeking behavior by inhibiting glutamate release and excitation of DA neurons in the NAc. In fact, human findings using positron emission tomography (PET) imaging with the selective mGluR5 radio tracer ^11^C-ABP688 reported no differences in mGluR5 density between chronic CU and healthy controls using [50], whereas other studies showed lower mGluR5 density in CU [51,52].

Only a few human studies have investigated endocannabinoids in biological samples of individuals with cocaine use so far. One study reported elevated plasma levels of AEA, and the structurally related N-acylethanolamines (NAEs) palmitoylethanolamide (PEA), and oleoylethanolamide (OEA) but lower 2-AG levels in abstinent individuals with cocaine use disorder (CUD) compared to healthy controls. Of note, reduced 2-AG levels in CUD was likely primarily driven by co-morbid alcohol use disorder (AUD) [53]. Moreover, we recently found lower levels of OEA and PEA, two NAEs that are linked to the metabolic pathway of AEA but do not bind directly at the CB_1_ receptor, in hair samples of chronic CU compared to stimulant-naïve healthy controls [54]. Given that hair analysis enables a retrospective determination of eCB/NAE concentrations over the last three months on average [55], our previous findings indicate tonic alterations in these lipid pathways in chronic CU. However, basal eCB/NAE plasma levels of current, chronic CU and its interaction with mGluR5 have yet to be studied in humans.

Here, we analyzed basal 2-AG, AEA, OEA, and PEA plasma levels of recreational and dependent CU and stimulant-naïve healthy controls at baseline and a 12-month follow-up to determine a specific targeted lipid profile potentially related to different levels of cocaine use in humans. Additionally, we used previously reported PET data of mGluR5 density of a subsample [50] to investigate potential cocaine-related associations between plasma eCB/NAE levels and mGluR5 density in an exploratory analysis. Based on reported animal model data and our recent findings [54], we expected alterations of eCB/NAE plasma levels primarily in dependent CU compared to controls, specifically for 2-AG, as well as an association between 2-AG and mGluR5 density in chronic CU.

## Methods and Materials

### Participants

As part of the longitudinal *Zurich Cocaine Cognition Study* [ZuCoSt; 56,57], plasma eCB/NAE data of stimulant-naiLve controls (n=92) and chronic CU (n=103; n=69 recreational and n=34 dependent users) were included. Additionally, we used eCB/NAE plasma data from the follow-up session after 12 months, including 43 stimulant-naiLve healthy controls and 33 chronic CU from the first session to check for stability and reliability of eCB/NAE plasma levels over time. Chronic CU who participated in the follow-up assessment were further subdivided into individuals with increased (n=16) and decreased (n=17) cocaine use. Inclusion and exclusion criteria are described in the supplementary materials. Cocaine dependence was diagnosed following the DSM-IV criteria [58], with dependent CU fulfilling and recreational CU not meeting these criteria.

Eighteen male chronic CU and 17 cocaine-naiLve controls from the *ZuCo^2^St* further underwent an additional PET procedure on a separate testing day to determine mGluR5 density (see below) with the same in-and exclusion criteria. Of this subsample, PET data of 18 male CU and 16 male controls were available with corresponding basal eCB/NAE plasma data.

The study has been carried out in accordance with the Declaration of Helsinki and was approved by the Cantonal Ethics Committee of Zurich (2022-00465, E-14/2009, 2009-0099/3). All participants provided written informed consent and were financially compensated for their participation.

### Analysis of N-acylethanolamines and phyto-cannabinoids in plasma

Blood plasma samples were taken after the psychiatric interviews. Four NAEs including the two eCBs, 2-AG and AEA, as well as OEA, and PEA and the plant-derived cannabinoids Δ^9^-tetrahydrocannabinol (THC) and cannabidiol (CBD) were extracted by liquid-liquid extraction and analyzed by isotope-dilution liquid chromatography/tandem mass spectrometry as previously described [59] using an Agilent 1200 high-performance liquid chromatography system coupled with an API 5000 triple quadruple mass spectrometer (AB Sciex).

### PET Imaging and Analysis

PET imaging acquisition and preprocessing have been previously described in detail [50]. Briefly, data were acquired with the whole-body scanner Discovery STX PET/computed tomography scanner (GE Healthcare, Waukesha, USA) at the Department of Nuclear Medicine of the University Hospital Zurich. The mGluR5-selective radioligand ^11^C-ABP688 [60] was administered according to a bolus-infusion-protocol [61].

PET images were preprocessed using PMOD 3.307 software (PMOD Technologies, Zurich, Switzerland). Nine VOIs were generated based on the Montreal Neurological Institute (MNI) brain templates: *Prefrontal Cortex (PFC)*, *hippocampal regions, anterior cingulate cortex (ACC), midcingulate cortex (MCC), amygdala, insula, caudate, putamen, and thalamus VOI*, as well as the cerebellar reference VOI. Normalized values of distribution (V_norm_=C_T[VOI]_/C_T[Cer]_) of ^11^C-ABP688 uptake were calculated [62]. For more information see supplementary materials.

All procedures were kept consistent on the baseline, follow-up, and PET testing days, when blood samples were taken (see supplementary materials).

### Statistical Analysis

Statistical analyses were performed with IBM SPSS Statistics 28.0.1.1. and R version 4.2.1 [63] for Tukey HSD post-hoc analyses using *multcomp* R package. Pearson’s χ^2^-tests were carried out to analyze frequency data and quantitative between-group data were analyzed by analyses of variance (ANOVAs) with *GROUP* (controls, recreational and dependent CU) as the fixed factor.

For the main outcome variables of eCB/NAE plasma concentrations, we used analyses of covariance (ANCOVAs) with *GROUP* as the fixed factor to control for the confounding variables sex [64], age [65], and recent cannabis use, i.e., THC and CBD plasma levels as well as current alcohol dependence based on their previously reported influences on 2-AG [53,66]. Repeated measure ANCOVA with sex and age as covariates and *TIME* (baseline and follow-up) as within-subject factor, and *GROUP* (controls, CU increase, and CU decrease) as between-subject factor as well as correlation analyses were used for a reliability check of eCB/NAE.

Potential associations between eCBs/NAEs and cocaine use variables (see Table 1) within chronic CU were investigated using stepwise linear regression analyses – *forward* and *backward* method (see supplementary materials). The influence of current alcohol dependence was specifically considered here given previously reported impact of co-morbid AUD on the association of 2-AG with CUD [53].

**Table 1.** Demographic data (means and standard deviations)

|  | **Stimulant-naïve controls**<br>n=92 | **Recreational cocaine users**<br>n=69 | **Dependent cocaine users**<br>n=34 | **Value** | **df1, df2** | **p** |
| --- | --- | --- | --- | --- | --- | --- |
| Female/male | 26/66 | 18/51 | 9/25 | $\chi^2 = 0.1$ | 2 | 0.949 |
| Age | 30.55 (9.1) | 28.67 (7.6) | 33.97 (10.6)<sup>†</sup> | F = 4.1 | 2, 192 | **0.018** |
| Years of education | 10.80 (1.8) | 10.38 (1.7) | 9.49 (1.2)<sup>°†</sup> | F = 7.9 | 2, 192 | **<0.001** |
| Verbal IQ | 108.02 (12.1) | 103.13 (9.7)<sup>°</sup> | 100.91 (11.4)<sup>°</sup> | F = 6.6 | 2, 192 | **0.002** |
| BDI sum score | 4.16 (4.2) | 7.67 (6.6)<sup>°</sup> | 11.53 (8.2)<sup>°†</sup> | F = 20.4 | 2, 192 | **<0.001** |
| ADHD-SR | 7.51 (4.9) | 12.90 (9.1)<sup>°</sup> | 16.88 (8.4)<sup>°†</sup> | F = 24.1 | 2, 192 | **<0.001** |
| Smoking (y/n) | 66/26 | 57/12 | 27/7 | $\chi^2 = 2.8$ | 2 | 0.251 |
| Cigarettes/week<sup>a</sup> | 84.14 (61.7) | 107.87 (55.9) | 135.04 (95.7)<sup>°</sup> | F = 5.8 | 2, 147 | **0.004** |
| Alcohol grams/week | 107.1 (115.6) | 170.70 (114.8) | 226.23 (322.0)<sup>°</sup> | F = 6.9 | 2, 192 | **0.001** |
| Alcohol dependence (y/n)<sup>c</sup> | - | 3/66 | 7/27 | $\chi^2 = 6.9$ | 1 | **0.009** |
| **Cocaine** |  |  |  |  |  |  |
| Cocaine craving | - | 19.14 (9.2) | 20.50 (11.4) | F = 0.4 | 1, 101 | 0.518 |
| Grams/week | - | 1.09 (1.4) | 5.47 (6.9) | F = 25.7 | 1, 101 | **<0.001** |
| Cocaine abstinence in h | - | 643.40 (881.0) | 535.07 (814.6) | F=0.4 | 1, 101 | 0.549 |
| Years of use<sup>b</sup> | - | 6 (0.75 - 17) | 9 (1 - 30) | F = 8.7 | 1, 101 | **0.004** |
| Hair concentration pg/mg | - | 3 070<br>(6 685.8) | 18 685.74<br>(21 478.2) | F = 30.7 | 1, 101 | **<0.001** |
| Urine sample positive (y/n) | - | 9/59 | 13/21 | $\chi^2 = 8.4$ | 1 | **0.004** |
| **Cannabis** |  |  |  |  |  |  |
| Cannabis use (y/n) | 40/52 | 46/23<sup>°</sup> | 22/12<sup>°</sup> | $\chi^2 = 10.0$ | 2 | **0.007** |
| Cannabis dependence (y/n)<sup>c</sup> | 3/89 | 6/63 | 3/31 | $\chi^2 = 2.52$ | 2 | 0.283 |
| Grams per week<sup>a</sup> | 1.21 (1.9) | 1.43 (2.4) | 2.49 (5.4) | F = 1.3 | 2, 105 | 0.286 |
| Years of use<sup>a,b</sup> | 7 (0 - 35) | 9 (0 - 26) | 12.5 (1.5 - 34)<sup>°†</sup> | F = 6.2 | 2, 105 | **0.003** |
| Urine sample positive (y/n)<sup>a</sup> | 12/28 | 14/31 | 11/11 | $\chi^2 = 2.9$ | 2 | 0.232 |
| **MDMA** |  |  |  |  |  |  |
| MDMA use (y/n) | 1/91 | 22/47<sup>°</sup> | 10/24<sup>°</sup> | $\chi^2 = 31.2$ | 2 | **<0.001** |
| Pills per week<sup>a</sup> | - | 0.29 (0.4) | 1.22 (3.1) | F = 1.8 | 1, 28 | 0.186 |
| Years of use<sup>a,b</sup> | - | 4.5 (0 - 17) | 5 (0.5 - 18) | F = 1.1 | 1, 28 | 0.294 |
| Hair concentration pg/mg<sup>a</sup> | - | 1 199.75<br>(2 336.5) | 673.00<br>(1 026) | F = 0.5 | 1, 28 | 0.504 |

Significant *p*-values are marked in bold<sup>°</sup>Tukey HSD post-hoc tests significant differences compared to controls<sup>†</sup>Tukey HSD post-hoc tests significant differences compared to RCU<sup>a</sup> Within users<sup>b</sup> Median (range)<sup>c</sup> Based on DSM-IV criteria
ADHD: attention-deficit/hyperactivity disorder, BDI: Beck's Depression Inventory, MDMA: 3,4 methylenedioxymethamphetamine

For our secondary outcome, to determine associations between eCBs/NAEs and mGluR5 density, we used Spearman’s rank correlations within the chronic CU and control group separately and the Fisher’s r-to-z transformation to compare correlation coefficients, as well as binary logistic regression models (see supplementary materials).

False discovery rate (FDR) was applied to account for multiple comparisons when needed [67]. Cohen’s *d* effect size was calculated by the means and pooled standard deviations of the groups [68].

All plasma eCB/NAE levels were winsorized in order to account for outliers based on Wilcox [69] (see supplementary materials).

The statistical comparisons were carried out with a significance level of *p*<.05 (two-tailed).

## Results

Participants’ demographic and substance use data are shown in Table 1.

### Dependent CU show elevated 2-AG plasma levels

ANCOVAs including sex, age, recent cannabis use, and current alcohol dependence as covariates and the dependent variables AEA, 2-AG, OEA, and PEA yielded a significant group effect only for 2-AG (F(2,187)=4.3, *p*=.014, *η_p_ ^2^*=.04) showing highest basal 2-AG plasma levels in dependent CU. Tukey HSD post-hoc comparisons identified significant elevated basal 2-AG plasma concentration in dependent CU compared to controls (*p*=.011, *d*=0.61) and to recreational CU (*p*=.042, *d*=0.52). In contrast, controls and recreational CU did not differ in 2-AG plasma levels (*p*=.825). No further group differences were found for AEA, PEA, and OEA (*p’s*>.165) as shown in Figure 1b-d.

**Figure 1.**
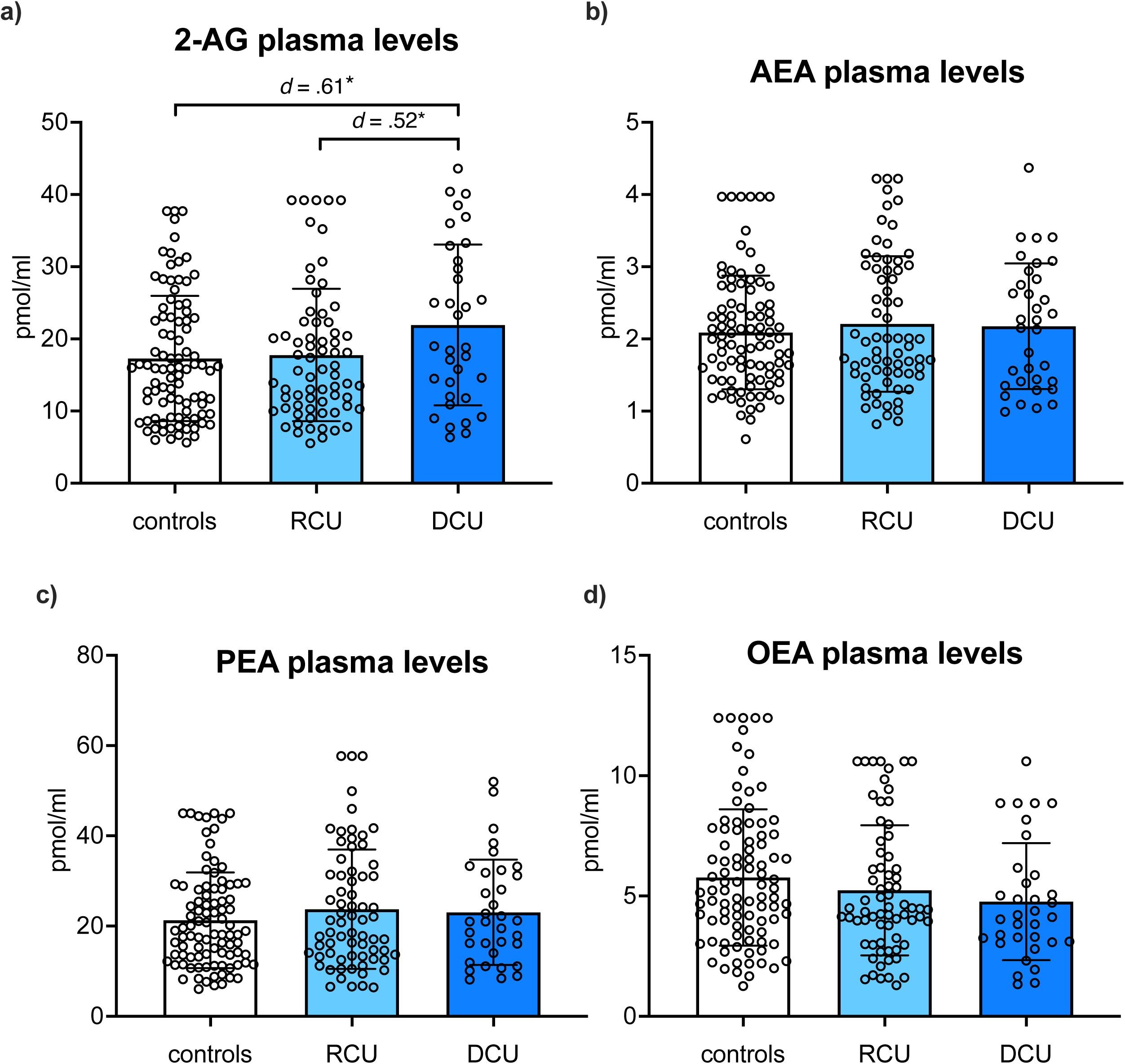
Dependent CU (DCU; dark blue) showed elevated basal plasma levels of **a)** 2-arachidonylglycerol (2-AG) compared to controls (white) and recreational CU (RCU; light blue), whereas no group differences were found for **b)** anandamide (AEA) **c)** palmitoylethanolamide (PEA) **d)** oleoylethanolamide (OEA). Bars represent means including individual data points, and error bars reflect standard error of the mean (SEM); corrected for age, sex, recent cannabis use, and alcohol dependence. Significant *p*-values marked with *p* < .05*, *p* < .01**

### 2-AG plasma concentration was stable and reliable over time

Repeated measure ANCOVA with the dependent variable 2-AG controlling for age and sex showed no main effect of *TIME* (F(1,72)=.03, *p*=.870) as well as no *TIME* x *GROUP* interaction (F(2,72)=.051, *p*=.950). All three groups showed stable 2-AG levels over the twelve months period even if cocaine use was increased or decreased (see supplementary materials Figure S1). Additionally, test-retest reliability was tested by correlation analyses with eCBs/NAEs plasma levels at baseline and follow-up (see supplementary materials Table S2). 2-AG plasma concentration at both time points was positively correlated over all participants and remained significant even after FDR-correction (*p*_FDR_=.016).

### Elevated basal 2-AG levels are associated with higher cocaine craving

Stepwise linear regression analyses *forward* and *backward* within chronic CU for each eCB/NAE (AEA, 2-AG, OEA, and PEA) as the dependent variable and entering cocaine use variables (see Table 1) as regressors yielded exactly one significant association. Over all four regression models regarding each eCB/NAE, only cocaine craving questionnaire (CCQ) sum score was identified as a significant regressor for basal 2-AG plasma level (F(1, 98)=7.1, *p*=0.009) even when controlling for age, sex, and recent cannabis use in the model (see supplementary materials Table S3). Separate analyses of recreational and dependent CU groups indicate that the negative association between 2-AG and CCQ sum score was driven by dependent CU (B=-0.41, t=-2.60, *p*=0.015), explaining 38% of the variance of basal 2-AG plasma levels (F(5, 28)=3.4, *p*=0.015, *R^2^*=0.38), in contrast to recreational CU (B=-0.13, t=-1.05, *p*=0.297). Association within recreational and dependent CU are shown in the Supplementary Figure S2a, and regression parameters of the models are shown in Supplementary Table S3.

To formally test whether current alcohol dependence and its interaction with 2-AG might influence the relationship between cocaine craving and 2-AG in dependent CU, we used a Linear Mixed Model (LMM) with *CCQ sum score* as the dependent variable and *2-AG*, *alcohol dependence*, and the interaction *2-AG*alcohol dependence* as factors. Results showed that the initial relationship between cocaine craving and 2-AG was primarily driven by alcohol dependence (F(1,34)=11.36, *p*=.002) and its interaction with 2-AG (F(1,34)=8.03, *p*=.008), whereas 2-AG alone was no longer a significant predictor for CCQ sum score in the model (F(1,34)=2.62, *p*=.115). More precisely, results indicate that the association between cocaine craving and 2-AG was only present in dependent CU co-morbid with alcohol dependence (see supplementary materials Figure S2b). Additional LMM including *cannabis dependence* in the model was not significant (see supplementary materials Table S4).

### Interaction between 2-AG and mGluR5 is specific to chronic cocaine use

Correlation analyses yielded significant positive correlations between 2-AG and region-specific mGluR5 VOIs within chronic CU showing significantly different correlation coefficients compared to controls for the brain structures *insula, ACC, MCC, amygdala, thalamus, caudate,* and *hippocampal regions* (see Table 2). Positive interactions within chronic CU are shown in Figure 2a-g (scatterplots for the controls are shown in the supplementary materials Figures S3). After controlling for multiple comparisons using FDR, only the correlation *p*-value of the thalamus region remained significant (see Table 2) while the other regions did not reach the significant threshold anymore even though showing clear statistical trends. To increase test power and to test for mGluR5 brain density overall, we performed an additional LMM including the fixed factors *2-AG* and *GROUP* as well as its interaction *2-AG*\**GROUP* and all nine cortical and subcortical mGluR5 VOIs for each subject as the dependent variable, with *subjects* and *mGluR5 VOIs* as random effect to control for subject and repeated level (VOIs within each subject) variability. LMM result showed a specific interaction effect of *2-AG*GROUP* on mGluR5 density overall (F(1,34)=6.33, *p*=.017), whereas *2-AG* alone (F(1,34)=1.29, *p*=.265) and *GROUP* (F(1,34)=4.07, *p*=.052) did not reach the significant threshold. The *2-AG*GROUP* interaction effect on mGluR5 remained significant even after including *smoking* and *age* into the model (F(1,34)=4.41, *p*=.043) indicating a robust positive association between 2-AG and mGluR5 brain density overall only in the CU group (see Figure 3). The model further yielded a significant main effect of *smoking* on mGluR5 overall density (F(1,34)=5.97, *p*=.020) but not *GROUP* alone (F(1,34)=2.63, *p*=.114) as we reported previously [50].

**Table 2.** Spearman’s correlation coefficients for 2-AG plasma levels and mGluR5 density.

|  | mGluR5 density VOIs | r | p | Fisher's z |
| --- | --- | --- | --- | --- |
| Cocaine users | Amygdala | **0.51** | **.033** | 2.39 |
| Controls | | -0.34 | .204 | $p=.017$ |
| Cocaine users | Insula | **0.52** | **.027** | 2.21 |
| Controls | | -0.26 | .339 | $p=.027$ |
| Cocaine users | ACC | **0.48** | **.046** | 2.12 |
| Controls | | -0.28 | .295 | $p=.034$ |
| Cocaine users | MCC | **0.48** | **.043** | 1.76 |
| Controls | | -0.14 | .602 | $p=.078$ |
| Cocaine users | Thalamus | **0.59** | **.011<sup>†</sup>** | 2.62 |
| Controls | | -0.31 | .240 | $p=.009†$ |
| Cocaine users | Caudate | **0.56** | **.016** | 2.56 |
| Controls | | -0.33 | .217 | $p=.011†$ |
| Cocaine users | Putamen | 0.42 | .083 | 1.91 |
| Controls | | -0.27 | .322 | $p=.056$ |
| Cocaine users | Hippocampal region | **0.47** | **.048** | 2.16 |
| Controls | | -0.30 | .264 | $p=.031$ |
| Cocaine users | PFC | 0.44 | .067 | 2.01 |
| Controls | | -0.28 | .295 | $p=.044$ |

Significant $p$ -values $<.05$ are shown in bold (uncorrected), <sup>†</sup>Significant after FDR-correction.

**Figure 2.**
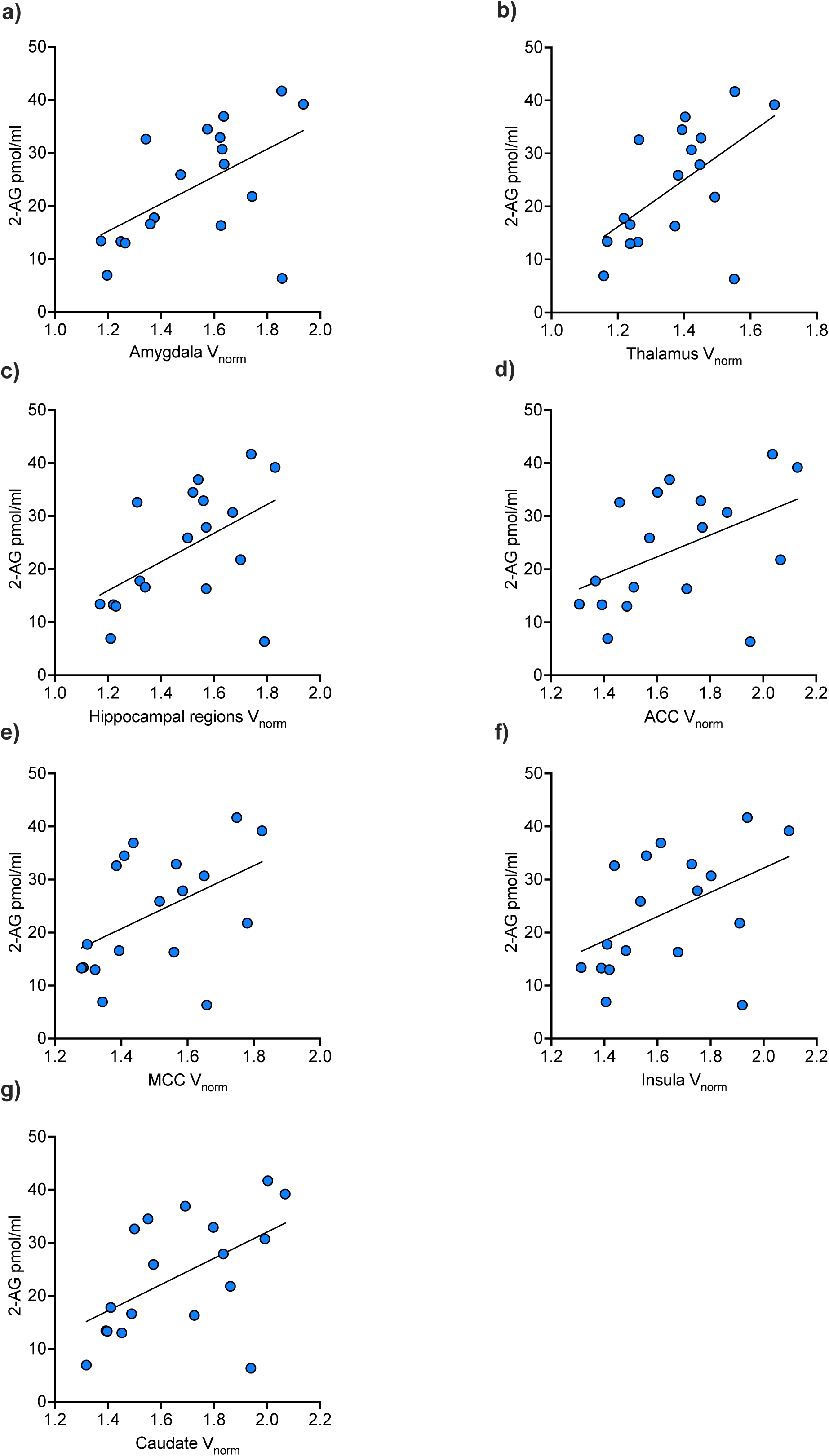
Within chronic cocaine users (n=18), basal plasma levels of 2-AG were positively associated with mGluR5 density of the **a)** amygdala, **b)** thalamus, **c)** hippocampal regions, **d)** anterior cingulate cortex (ACC), **e)** midcingulate cortex (MCC) **f)** insula **g)** caudate. After controlling for tobacco use and age, the interaction between 2-AG and mGluR5 was specifically evident in chronic cocaine users for the brain structures amygdala **(a),** thalamus **(b)**, as well as hippocampal region **(c)**, and as trend levels for the other brain structures (d-g).

**Figure 3.**
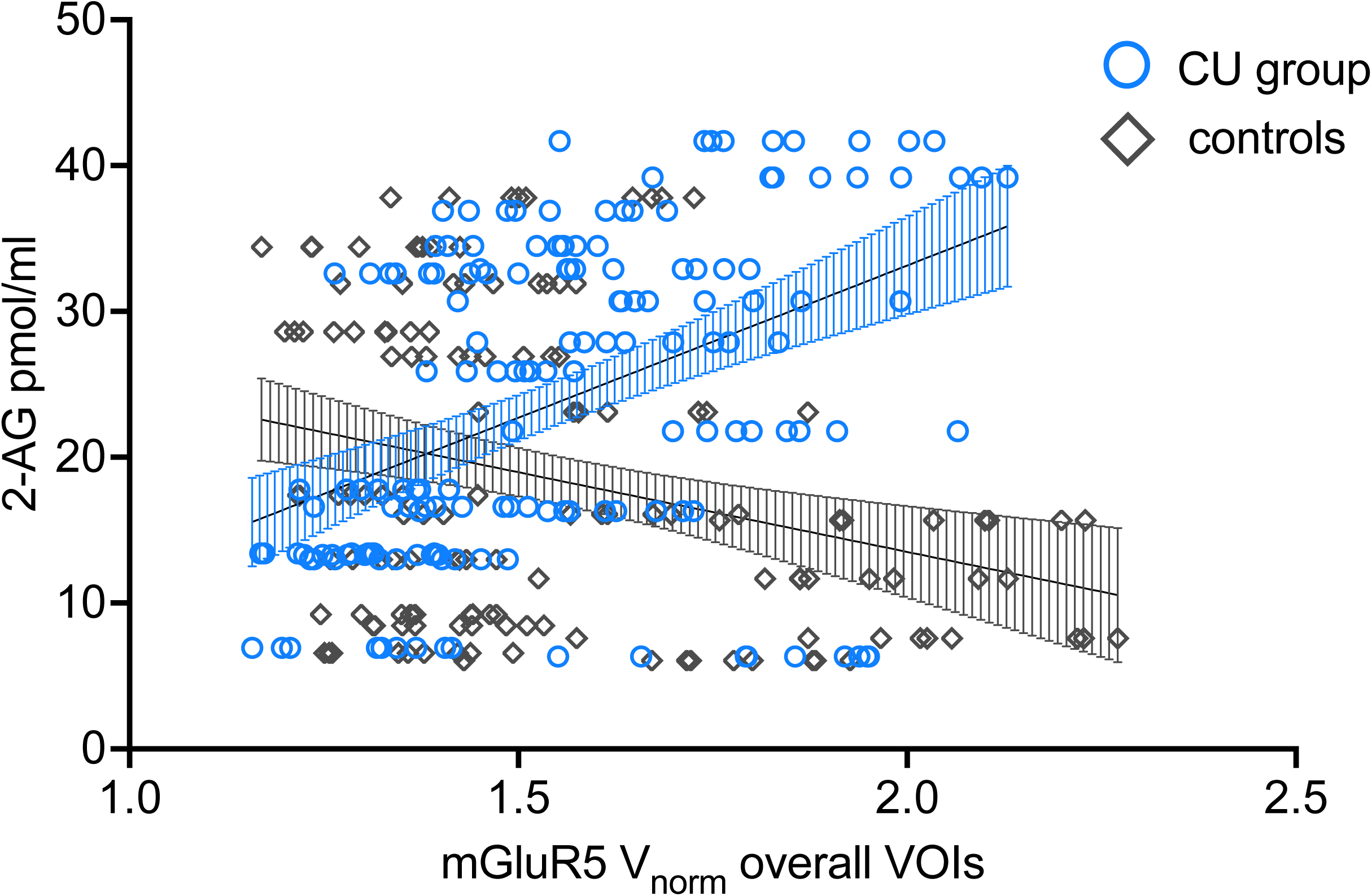
Linear mixed model yielded a specific interaction effect of *2-AG*GROUP* on mGluR5 density overall indicating a positive association between 2-AG and mGluR5 density overall VOIs only in chronic cocaine users (CU; blue circles) but not in controls (gray diamonds).

To formally address whether the relationship between 2-AG and mGluR5 density was unique to chronic CU even after controlling for confounding variables age and smoking, we used binary logistic regression models with *GROUP* as the dependent variable. Robust interaction effects for chronic CU between 2-AG and mGluR5 density were found for the brain structures: *thalamus* (*B*=1.09, Wald=4.61, *p*=.032, OR=2.96, 95%CI [1.10; 7.98]), *amygdala* (*B*=0.62, Wald=4.34, *p*=.037, OR=1.85, 95%CI [1.04; 3.30]), and *hippocampal regions* (*B*=0.77, Wald=4.39, *p*=.036, OR=2.15, 95%CI [1.05; 4.40]). *Insula, caudate, ACC,* and *MCC* showed differences at trend levels but did not meet the significant threshold (see supplementary materials Table S5d-g). All logistic regression models are shown in the supplementary materials Table S5.

## Discussion

Our findings indicate alterations of the ECS in individuals with cocaine dependence. Specifically, basal 2-AG was elevated in cocaine dependence compared to controls and at a trend level also to recreational users. In contrast, the latter two groups did not differ in basal 2-AG plasma concentration. Additional exploratory analyses of a male subgroup with mGluR5-specific PET yielded a positive association between basal 2-AG plasma levels and cerebral mGluR5 density only evident in chronic CU, corroborating animal findings and suggesting a specific 2-AG/mGluR5 interaction underpinning cocaine dependence also in humans.

Our findings suggest that 2-AG might play a crucial role in cocaine dependence in humans. This is consistent with recent animal models showing elevated 2-AG levels in the VTA after cocaine administration, contributing to the positive reinforcing and motivation-enhancing effects of the substance [40,43,70]. Although human plasma samples can only reflect peripheral eCB/NAE levels and do not allow direct assessment of brain-specific concentrations, we speculate that the elevated 2-AG levels found in individuals with cocaine dependence indicate higher response to the cocaine-rewarding effects as well as higher motivated behavior resulting in increased vulnerability to developing cocaine dependence. Given that i) cocaine use intensity (i.e., cocaine grams per week, cocaine years of use, cocaine abstinence, cocaine hair concentrations, and positive urine sample) did not significantly explain the 2-AG variance, ii) 2-AG levels were stable over time, and iii) 2-AG levels did not covary with changing cocaine use between baseline and follow-up, elevated basal 2-AG plasma levels might reflect a risk factor for cocaine dependence (e.g., reward behavior) rather than a dose-dependent consequence of chronic cocaine use. Accordingly, no correlations between 2-AG and cocaine addiction severity variables have been previously shown in abstinent cocaine users [53]. In contrast to our findings, the same study found reduced 2-AG plasma levels and elevated NAE levels in abstinent individuals with CUD compared to healthy controls. However, lower 2-AG plasma levels were only significantly different from controls in abstinent cocaine users with co-morbid AUD as the authors reported in the supplements, whereas AUD had no influence on NAE levels. Moreover, a high number of psychiatric co-morbidities such as mood (40%), anxiety (25%) and substance use disorder other than cocaine (86%) as well as personality disorders (33%) were reported for the abstinent CUD group, which had effects on eCB/NAE plasma levels and might further explain the different findings of 2-AG. Importantly, ROC curves showed that CUD was only a significant predictor for NAEs but not 2-AG indicating that the findings regarding lower 2-AG plasma levels in abstinent cocaine users might not be related to chronic cocaine use per se. Since we controlled for a variety of confounding variables, which have been reported to influence eCB/NAE, we believe that our analyses detected robust and specific effects of dependent cocaine use. However, the question of whether elevated 2-AG levels in individuals with cocaine dependence is due to long-term changes in synaptic plasticity caused by repeated cocaine use, as previously shown in animal models, or whether it is a pre-existing phenomenon, and 2-AG is thus a biomarker of vulnerability to cocaine dependence, requires further investigation. Our present analyses with follow-up data suggest stable 2-AG plasma levels over a period of at least 12 months, independent of increased or decreased cocaine use, which might be indicative of 2-AG being a biomarker (i.e., trait marker) or alternatively suggesting that cocaine-induced changes in 2-AG might be of limited reversibility. Future studies should therefore address our assumption that individuals with elevated basal 2-AG levels are more prone to the rewarding effects of cocaine (e.g., wanting and liking) and subsequently more vulnerable to developing cocaine dependence.

Although we found higher basal 2-AG plasma levels in the cocaine-dependent group, cocaine craving was negatively associated with 2-AG plasma levels in dependent CU initially. This result would have been in contrast to previous animal findings reporting a positive association between stress-induced cocaine craving and phasic 2-AG response [71,72]. However, the additional LMM showed that the relationship between 2-AG and cocaine craving was primarily driven by alcohol dependence and only found in individuals with co-morbid cocaine and alcohol dependence but not with cocaine dependence alone. Genetic and pharmacological downregulation of 2-AG signaling in rodent brain regions responsible for stress and affect regulation (e.g., amygdala, paraventricular hypothalamus, and prefrontal regions) has been linked to anxiety-and depressive-like effects typically observed during drug withdrawal [73,74]. Therefore, one might speculate that our findings of lower 2-AG levels associated with higher cocaine craving in individuals with co-morbid cocaine and alcohol dependence may be a phasic response to the negative affective state of cocaine and/or alcohol withdrawal. However, no reliable measures of withdrawal and anxiety symptoms neither for cocaine nor for alcohol were available for the present dataset and future human studies should investigate the influence of alcohol and cocaine withdrawal on 2-AG and craving in AUD alone and co-morbid with CUD. In fact, since the relationship between 2-AG and cocaine craving was no longer evident in dependent CU without co-morbid alcohol dependence, the present results support our suggestion of elevated 2-AG levels as a stable biomarker unique for cocaine dependence.

Here, we found for the first time a relationship between 2-AG and mGluR5 density in humans, including brain regions involved in processing reward behavior and affect regulation (i.e., amygdala, thalamus, and hippocampal regions). This association was evident only in chronic CU but not in healthy controls and strongest in the thalamus, where the correlation survived FDR-correction despite the small sample size. Moreover, the LMM including all nine mGluR5 density VOIs yielded a robust effect of 2-AG on mGluR5 density overall specific for chronic CU. Our present results confirmed previous animal models showing an involvement of mGluR5 and 2-AG in LTD throughout various brain regions [45-47] and are consistent with the notion of drug-induced long-term synaptic plasticity [1]. Notably, single *in vivo* cocaine administration has been previously reported to affect mGluR5 and endocannabinoid-LTD in mice [44], which might further explain that our finding of the statistical interaction between 2-AG and mGluR5 was only evident in CU. Recent animal models reported an association between the mGluR5-mediated 2-AG-LTD in the NAc and reward-seeking behavior, showing that drug-conditioned cues can amplify 2-AG-mediated LTD by excitatory afferents from the PFC impinging on D1-MSN in the NAc and stimulating glutamate release [25,49]. Subsequent activation of postsynaptic mGluR5-mediated 2-AG-LTD amplified LTD at glutamatergic neurons resulting in increased drug-seeking behavior, including craving. These findings are consistent with both previous preclinical results and recent human findings reporting impaired glutamate homeostasis in the NAc in chronic CU [75,76]. Although we were not able to assess mGluR5 density in the ventral striatum (i.e., VTA and NAc) due to limited spatial resolution of PET imaging, and eCBs/NAEs can only be assessed in the periphery and not brain-structure-specific in humans, we nevertheless found a significant interaction between basal 2-AG and mGluR5 density in a variety of brain regions involved in reward and affect regulation, which was evident only in chronic CU. These findings indeed indicate that there might be a specific 2-AG/mGluR5-mechanism underpinning cocaine dependence in humans, which may contribute to increased craving behavior and vulnerability to cocaine relapse. Our results extend previous findings from our lab [50], indicating that cocaine use disorder might be rather linked to 2-AG and its interaction with mGluR5 than exclusively to mGluR5 density per se, which has been reported previously [51,52]. Of note, our PET subsamples consist of male controls and CU and need to be tested in women as well. Future studies should address this issue and our assumption by using PET imaging techniques with selective tracers for enzymes and lipids related to 2-AG in both sexes. This will allow for detecting alterations of the ECS, specifically in the NAc, during craving.

We recently reported lower tonic OEA and PEA hair concentration in a subgroup of the present chronic CU compared to healthy controls, specifically pronounced in individuals showing cocaine dependence [54]. However, we did not find differences between groups in AEA and 2-AG hair concentrations. This might be caused by the fact that concentrations of OEA and PEA are generally higher than AEA and 2-AG in hair and other biological samples [77]. In addition, both OEA and PEA, although structurally related to AEA, serve different functions in the body. While OEA is a satiety compound and the main endogenous ligand to PPAR-α and thus deeply involved in the regulation of the energy metabolism [78], PEA is also considered an agonist at PPAR-α but also binds to the cannabinoid-like G-protein coupled receptors GPR55 and GPR119 and is suggested to be related to chronic inflammation and pain [79,80]. Moreover, reported findings might also be related to phasic vs. tonic eCB/NAE release in the brain, which may be further supported by our null results of the correlation analyses between plasma and hair levels of eCBs/NAEs after FDR-correction (see supplementary materials Table S6). No association between hair and plasma eCB/NAE levels have been also recently reported [81].

The present findings are novel in the field of cocaine use disorder in humans and crucially contribute to a better mechanistic understanding of the role of the ECS in cocaine dependence. Specifically, 2-AG seems to be a critical player and might be a promising pharmaco-therapeutic target for novel treatments of cocaine use disorder aiming to prevent drug relapse and improve cocaine abstinence.

## Acknowledgment

We thank Avery Nelson for his preliminary statistical analyses with parts of the present data set.

## Author contributions

SLK and BBQ contributed to the final research question and hypotheses. SLK analyzed the data and wrote the paper. BBQ designed initial research and revised the paper. LMH, MV, and KHP were involved in data collection and AKK helped with the dataset. VT and SMA were involved in the PET study and supported initial PET data analysis. MRB performed the toxicological hair analysis. CB, FP, CR, and FML conducted the eCB/NAE plasma analyses.

