## Supplementary Materials for "Plasma endocannabinoids in cocaine dependence and their interaction with cocaine craving and metabotropic glutamate receptor 5 density in the human brain"

**Methods and Materials**

**Participants**

Blood samples of, in total, 224 individuals were collected as part of the longitudinal *Zurich Cocaine Cognition Study* [ZuCoSt; 1,2] of which eCB/NAE plasma levels of 195 individuals were available for present analyses.

Inclusion criteria for chronic CU were cocaine use of at least 1 g per month, cocaine as the primary used illegal substance, and a current abstinence duration <6 months. Exclusion criteria for the user groups were past or current use of opioids, a polysubstance use pattern, and an axis-I DSM-IV adult psychiatric disorder except for cocaine, cannabis, tobacco, and alcohol abuse/dependence, as well as a history of affective disorders and attention deficit hyperactivity disorder (ADHD). Exclusion criteria for the control group were acute or history of any axis-I DSM-IV psychiatric disorder and any form of addiction or regular substance use (lifetime use <15 occasions), with exception of cannabis and alcohol. All participants were asked to abstain from illegal substances for a minimum of 72 h and not to consume alcohol for a minimum of 24 h before the test session. Abstinence, and self-reported substance use was controlled via urine and 6-months hair analysis, respectively.

**PET Imaging and Analysis**

The mGluR5-selective radioligand ^11^C-ABP688 [3] was administered according to a bolus-infusion-protocol [4], including a bolus-administration of half of the tracer over 2 minutes followed by tracer-infusion over 58 minutes of the other half.

Radioactivity of ^11^C-ABP688 was measured for 60 minutes using 20 frames (10x60s, 10x300s). Prior to the PET acquisition, T1-weighted magnetic resonance imaging was performed on a Philips Achieva 3T scanner (Philips Healthcare, Best, The Netherlands) for PET imaging preprocessing.

Frames 17-19 were averaged, co-registered to the individual MR image and spatially normalized to the Montreal Neurological Institute (MNI) space via segmentation and normalization of the individual T1-weighted anatomical images. Volumes of interested (VOIs) were derived from the segmented AAL template (excluding white matter portions), which is based on the MNI template. Frontal VOIs were merged into one *prefrontal cortex (PFC)* VOI, including dorsolateral PFC, medial PFC, ventrolateral PFC, and orbitofrontal cortex. Moreover, we combined the VOIs hippocampus and parahippocampal gyrus into *hippocampal regions*.

Normalized values of distribution (V_norm_=C_T[VOI]_/C_T[Cer]_) of ^11^C-ABP688 uptake were calculated by dividing the average radioactivity concentration between 45 and 55 minutes (frames 17-19) of each VOI (C_T[VOI]_) with the corresponding cerebellar radioactivity concentration (C_T[Cer]_), which is equivalent to BP_ND_ +1 [5].

**Procedure**

All participants were examined by trained psychologists using the Structural Clinical Interview for DSM-IV disorders SCID-I [6]. Substance use was determined by using the standardized and structured Interview for Psychotropic Drug Consumption [7] as well as objectively by urine, blood (for ∆^9^-tetrahydrocannabinol [THC] and cannabidiol [CBD], see below), and toxicological hair analysis [2]. Current cocaine craving was assessed by the brief version of the Cocaine Craving Questionnaire (CCQ) [8]. The severity of nicotine dependence was determined by the Fagerström Test of Nicotine dependence [9]. ADHD was assessed by the self-reports using the ADHD self-rating scale [10], and current symptoms of depression were measured by the Beck Depression Inventory (BDI) [11].

On average, test sessions between eCB/NAE plasma sampling and PET were separated by 137 days (median=79 days; range: -218–689 days).

**Statistical Analysis**

More precisely, cocaine use variables were entered as regressors into the model with basal plasma levels of AEA, 2-AG, OEA, and PEA as the dependent variables for each model. The *forward* and *backward* method was performed to account for suppressor effects as a validation check with a significance level of *p* <.05. We checked for multicollinearity by using the variance inflation factor (VIF) with a cutoff value >10 and the tolerance statistic with a cutoff value <0.1 [12,13]. None of the independent variables showed a violation of multicollinearity (see Table S1a-b).

To formally test whether results from the correlation analyses were unique to one of the groups even after controlling for the confounding variable age for eCBs/NAEs and smoking status for mGluR5 density [14], we used binary logistic regression models with *GROUP* as the dependent variable, dummy coded with 0=controls and 1=chronic CU, and entered significant mGluR5 VOIs, eCBs/NAEs, their interaction as well as the confounding variables age and smoking as fixed factors.

**Winsorized method for endocannabinoid plasma levels**

All values larger than the value corresponding to the 75th percentile plus 1.5 times the interquartile range per group were replaced with the maximum value within this range in the corresponding group. Accordingly, mean values smaller than 25th percentiles minus 1.5 times the interquartile range per group were replaced with the minimal value within this range in the corresponding group [15].

**Results**

**Elevated basal 2-AG levels are associated with higher cocaine craving in dependent CU, which is explained by current alcohol dependence**

To control for age, sex, and recent cannabis use, we used a post-hoc *forced entry* linear regression model with the dependent variable 2-AG and the regressors CCQ sum score as well as the control variables. This model explained 19% of the variance of 2-AG basal plasma levels within chronic CU (F(5, 97)=4.7, *p*<0.001, *R^2^*=0.19), indicating a decrease of basal 2-AG plasma level of about 0.22 pmol/ml for each increase of one score in the CCQ (see Table S3a).

After splitting the chronic CU group into dependent and recreational CU, CCQ sum score was not a significant regressor in recreational CU (B=-0.13, t=-1.05, *p*=0.297), and the model explained 19% of the variance of the dependent variable (F(5, 63)=3.0, *p*=0.019, *R^2^*=0.19).

**Table S1** Multicollinearity statistics for the dependent variable: 2-AG

a)

|  | Tolerance | VIF |
| --- | --- | --- |
| CCQ sum score | 1.00 | 1.00 |
| Cocaine grams per week | 1.00 | 1.00 |
| Cocaine years of use | 0.98 | 1.02 |
| Cocaine abstinence in h | 0.99 | 1.01 |
| Cocaine hair concentration | 1.00 | 1.00 |
| Cocaine urine | 1.00 | 1.00 |

b)

|  | Tolerance | VIF |
| --- | --- | --- |
| CCQ sum | 0.94 | 1.07 |
| Sex | 0.94 | 1.06 |
| Age | 0.91 | 1.10 |
| THC blood level | 0.90 | 1.12 |
| CBD blood level | 0.88 | 1.14 |

Multicollinearity statistics for a) stepwise and b) forced entry linear regression analyses using the variance inflation factor (VIF) with a cutoff value > 10 and the tolerance statistic with a cutoff value < 0.1

**Table S2** Spearman’s correlation coefficients for eCB/NAE plasma levels at baseline and follow-up over all and within controls (n=44), CU increase (n=16), and CU decrease (n=17)

|  | Follow-up | baseline  AEA | baseline  2-AG | baseline  OEA | baseline  PEA |
| --- | --- | --- | --- | --- | --- |
| Controls | AEA | 0.26 | -0.09 | 0.21 | 0.25 |
| Increaser |  | -0.15 | 0.13 | 0.09 | -0.05 |
| Decreaser |  | 0.51 | -0.10 | 0.08 | 0.11 |
| Over all |  | 0.27 | -0.09 | 0.15 | 0.16 |
| Controls | 2-AG | 0.17 | 0.35 | 0.19 | 0.03 |
| Increaser |  | -0.02 | 0.45 | -0.03 | -0.51 |
| Decreaser |  | 0.25 | 0.17 | 0.10 | 0.14 |
| Over all |  | 0.11 | **0.39** | 0.04 | -0.02 |
| Controls | OEA | 0.33 | -0.15 | 0.40 | 0.38 |
| Increaser |  | -0.17 | -0.21 | 0.55 | 0.04 |
| Decreaser |  | 0.53 | -0.12 | -0.09 | 0.29 |
| Over all |  | 0.28 | -0.18 | **0.33** | 0.27 |
| Controls | PEA | 0.19 | -0.19 | 0.19 | 0.23 |
| Increaser |  | -0.44 | -0.03 | 0.05 | -0.32 |
| Decreaser |  | 0.51 | -0.16 | -0.16 | 0.37 |
| Over all |  | 0.17 | -0.20 | 0.11 | 0.18 |

Significant FDR-corrected *p*-values <.05 are shown in bold

**Table S3** Regression model parameters for the dependent variable: 2-AG

a)

|  | *B* | SE *B* | *β* | | t-value | *p* |
| --- | --- | --- | --- | --- | --- | --- |
| Constant | 8.16 | 5.34 |  |  | 1.53 | 0.130 |
| CCQ sum | -0.22 | 0.10 | -0.22 |  | -2.31 | **0.023** |
| Sex | 3.98 | 2.13 | 0.18 |  | 1.87 | 0.064 |
| Age | 0.28 | 0.11 | 0.25 |  | 2.62 | **0.010** |
| THC blood level | 0.11 | 0.10 | 0.10 |  | 1.07 | 0.286 |
| CBD blood level | -6.64 | 3.10 | -0.21 |  | -2.14 | **0.035** |

b)

|  | *B* | SE *B* | *β* | | t-value | *p* |
| --- | --- | --- | --- | --- | --- | --- |
| Constant | 4.59 | 6.75 |  |  | 0.68 | 0.499 |
| CCQ sum | -0.13 | 0.12 | -0.24 |  | -1.05 | 0.297 |
| Sex | 6.85 | 2.43 | 0.33 |  | 2.81 | **0.007** |
| Age | 0.11 | 0.14 | 0.10 |  | 0.81 | 0.420 |
| THC blood level | 0.15 | 0.11 | 0.16 |  | 1.41 | 0.163 |
| CBD blood level | -8.51 | 39.67 | -0.03 |  | -0.22 | 0.831 |

c)

|  | *B* | SE *B* | *β* | | t-value | *p* |
| --- | --- | --- | --- | --- | --- | --- |
| Constant | 22.54 | 9.12 |  |  | 2.47 | 0.020 |
| CCQ sum | -0.41 | 0.16 | -0.42 |  | -2.60 | **0.015** |
| Sex | -2.02 | 3.97 | -0.08 |  | -0.51 | 0.615 |
| Age | 0.37 | 0.17 | 0.35 |  | 2.17 | **0.039** |
| THC blood level | 0.04 | 0.25 | 0.03 |  | 0.16 | 0.875 |
| CBD blood level | -7.64 | 4.02 | -0.37 |  | -1.90 | 0.068 |

Linear regression models *forced entry* a) over all chronic cocaine users (CU) and within b) recreational CU and c) dependent CU with significant regressors shown in bold.

**Table S4** Linear mixed model parameters for the dependent variable: CCQ sum score

|  | Estimate | SE | CI | df | t | *p* |
| --- | --- | --- | --- | --- | --- | --- |
| (Intercept) | 29.83 | 4.01 | 21.67 – 37.98 | 34 | 7.43 | <.001 |
| 2-AG | -0.42 | 0.16 | -0.75 – -0.09 | 34 | -2.62 | **.013** |
| Cannabis dependence | 16.88 | 21.83 | -27.47 – 61.24 | 34 | 0.77 | .445 |
| Cannabis dependence*2-AG | -1.02 | 1.17 | -3.40 – 1.36 | 34 | -0.88 | .388 |

**Table S5** Binary logistic regression models for the dependent variable *GROUP*

a)

|  | *B* | SE *B* | *Wald* | | *p* | | *OR* | *95% CI for OR* |
| --- | --- | --- | --- | --- | --- | --- | --- | --- |
| Constant | 19.64 | 11.24 |  |  | |  |  |  |
| 2-AG | -1.41 | 0.67 | 4.44 |  | | **0.035** | 0.24 | 0.07; 0.91 |
| Thalamus | -14.98 | 8.18 | 3.35 |  | | 0.067 | <.001 | 0.00; 2.90 |
| Thalamus x 2-AG | 1.09 | 0.51 | 4.61 |  | | **0.032** | 2.96 | 1.10; 7.98 |
| Age | -0.04 | 0.05 | 0.57 |  | | 0.452 | 0.96 | 0.86; 1.07 |
| Smoking | 1.22 | 1.51 | 0.65 |  | | 0.420 | 3.38 | 0.18; 64.75 |

b)

|  | *B* | SE *B* | *Wald* | | *p* | | *OR* | *95% CI for OR* |
| --- | --- | --- | --- | --- | --- | --- | --- | --- |
| Constant | 7.95 | 6.67 |  |  | |  |  |  |
| 2-AG | -0.87 | 0.44 | 3.96 |  | | **0.047** | 0.42 | 0.18; 0.99 |
| Amygdala | -5.96 | 4.17 | 2.04 |  | | 0.153 | .003 | 0.00; 9.19 |
| Amygdala x 2-AG | 0.62 | 0.30 | 4.34 |  | | **0.037** | 1.85 | 1.04; 3.30 |
| Age | -0.04 | 0.05 | 0.59 |  | | 0.442 | 0.96 | 0.87; 1.07 |
| Smoking | 1.98 | 1.59 | 1.57 |  | | 0.211 | 7.28 | 0.31; 162.58 |

c)

|  | *B* | SE *B* | *Wald* | | *p* | | *OR* | *95% CI for OR* |
| --- | --- | --- | --- | --- | --- | --- | --- | --- |
| Constant | 10.63 | 7.35 |  |  | |  |  |  |
| 2-AG | -1.05 | 0.51 | 4.19 |  | | **0.041** | 0.35 | 0.13; 0.96 |
| Hippocampal | -8.11 | 4.92 | 2.72 |  | | 0.099 | <.001 | 0.00; 4.63 |
| Hippocampal x 2-AG | 0.77 | 0.37 | 4.39 |  | | **0.036** | 2.15 | 1.05; 4.40 |
| Age | -0.04 | 0.05 | 0.63 |  | | 0.428 | 0.96 | 0.86; 1.07 |
| Smoking | 1.98 | 1.79 | 1.22 |  | | 0.269 | 7.21 | 0.22; 240.31 |

d)

|  | *B* | SE *B* | *Wald* | | *p* | | *OR* | *95% CI for OR* |
| --- | --- | --- | --- | --- | --- | --- | --- | --- |
| Constant | 9.43 | 7.37 |  |  | |  |  |  |
| 2-AG | -0.57 | 0.36 | 2.48 |  | | 0.115 | 0.56 | 0.28; 1.15 |
| Insula | -5.92 | 4.23 | 1.96 |  | | 0.162 | .003 | 0.00; 10.72 |
| Insula x 2-AG | 0.39 | 0.23 | 2.80 |  | | 0.094 | 1.47 | 0.94; 2.31 |
| Age | -0.03 | 0.05 | 0.35 |  | | 0.552 | 0.97 | 0.88; 1.07 |
| Smoking | 0.39 | 1.33 | 0.09 |  | | 0.769 | 1.48 | 0.11; 19.91 |

e)

|  | *B* | SE *B* | *Wald* | | *p* | | *OR* | *95% CI for OR* |
| --- | --- | --- | --- | --- | --- | --- | --- | --- |
| Constant | 9.91 | 7.11 |  |  | |  |  |  |
| 2-AG | -0.54 | 0.34 | 2.48 |  | | 0.115 | 0.58 | 0.30; 1.14 |
| ACC | -5.97 | 3.97 | 2.26 |  | | 0.133 | .003 | 0.00; 6.15 |
| ACC x 2-AG | 0.36 | 0.21 | 2.83 |  | | 0.093 | 1.43 | 0.94; 2.16 |
| Age | -0.03 | 0.05 | 0.36 |  | | 0.550 | 0.97 | 0.88; 1.07 |
| Smoking | 0.15 | 1.26 | 0.01 |  | | 0.909 | 1.16 | 0.10; 13.76 |

f)

|  | *B* | SE *B* | *Wald* | | *p* | | *OR* | *95% CI for OR* |
| --- | --- | --- | --- | --- | --- | --- | --- | --- |
| Constant | 15.34 | 8.89 |  |  | |  |  |  |
| 2-AG | -0.74 | 0.45 | 2.76 |  | | 0.097 | 0.48 | 0.20; 1.14 |
| MCC | -10.08 | 5.62 | 3.21 |  | | 0.073 | <.001 | 0.00; 2.56 |
| MCC x 2-AG | 0.53 | 0.31 | 3.00 |  | | 0.083 | 1.70 | 0.93; 3.10 |
| Age | -0.03 | 0.05 | 0.31 |  | | 0.575 | 0.97 | 0.88; 1.08 |
| Smoking | -0.33 | 1.22 | 0.07 |  | | 0.790 | 0.72 | 0.07; 7.92 |

g)

|  | *B* | SE *B* | *Wald* | | *p* | | *OR* | *95% CI for OR* |
| --- | --- | --- | --- | --- | --- | --- | --- | --- |
| Constant | 12.16 | 7.77 |  |  | |  |  |  |
| 2-AG | -0.77 | 0.41 | 3.45 |  | | 0.063 | 0.46 | 0.21; 1.04 |
| Caudate | -7.48 | 4.39 | 2.90 |  | | 0.088 | .001 | 0.00; 3.08 |
| Caudate x 2-AG | 0.50 | 0.26 | 3.70 |  | | 0.054 | 1.66 | 0.99; 2.76 |
| Age | -0.04 | 0.05 | 0.69 |  | | 0.408 | 0.96 | 0.86; 1.06 |
| Smoking | 0.75 | 1.61 | 0.22 |  | | 0.640 | 2.12 | 0.09; 49.75 |

Binary logistic regression models entering the brain structures a) thalamus b) amygdala c) hippocampal regions d) insula e) ACC f) MCC g) caudate with significant regressors shown in bold.

**Correlations between hair and plasma eCB/NAE levels**

We previously published hair concentrations of eCBs/NAEs from an overlapping sample [16]. Thus, we explored potential correlations between hair and plasma eCB/NAE concentrations across all groups at baseline (see Table S6). However, we found only weak negative correlations between plasma 2-AG, AEA, OEA and hair 2-AG, AEA, OEA, and PEA concentrations, which did not survive FDR-correction except for the negative correlation between plasma OEA and hair 2-AG in controls (see Table S6).

**Table S6** Spearman’s correlation coefficients for eCB/NAE levels in plasma and hair over all and within controls (n=64), Recreational CU (RCU n=44), and Dependent CU (DCU n=25).

|  | Hair | plasma  AEA | plasma  2-AG | plasma  OEA | plasma  PEA |
| --- | --- | --- | --- | --- | --- |
| Controls | AEA | -0.25 | -0.15 | -0.21 | -0.19 |
| RCU |  | -0.21 | -0.05 | -0.20 | -0.11 |
| DCU |  | 0.10 | -0.37 | 0.07 | -0.04 |
| Over all |  | -0.19 | -0.21 | -0.15 | -0.15 |
| Controls | 2-AG | -0.24 | -0.21 | **-0.37** | -0.17 |
| RCU |  | -0.34 | -0.06 | -0.22 | -0.23 |
| DCU |  | 0.15 | -0.23 | 0.21 | 0.34 |
| Over all |  | -0.18 | -0.17 | -0.22 | -0.10 |
| Controls | OEA | -0.20 | -0.14 | -0.22 | -0.16 |
| RCU |  | -0.11 | -0.32 | 0.13 | -0.16 |
| DCU |  | 0.01 | -0.40 | 0.21 | 0.07 |
| Over all |  | -0.15 | -0.25 | 0.004 | -0.14 |
| Controls | PEA | -0.21 | -0.04 | -0.13 | -0.22 |
| RCU |  | -0.16 | -0.29 | 0.15 | -0.11 |
| DCU |  | -0.01 | -0.44 | 0.25 | 0.02 |
| Over all |  | -0.18 | -0.22 | 0.03 | -0.16 |

Significant FDR-corrected *p*-values <.05 are shown in bold

**Figure S1** Repeated measure ANCOVA with *TIME* (baseline and follow-up) as within subject factor and *GROUP* (controls: black circle, CU increase: orange triangle, and CU decrease: green diamonds) as the between factor.

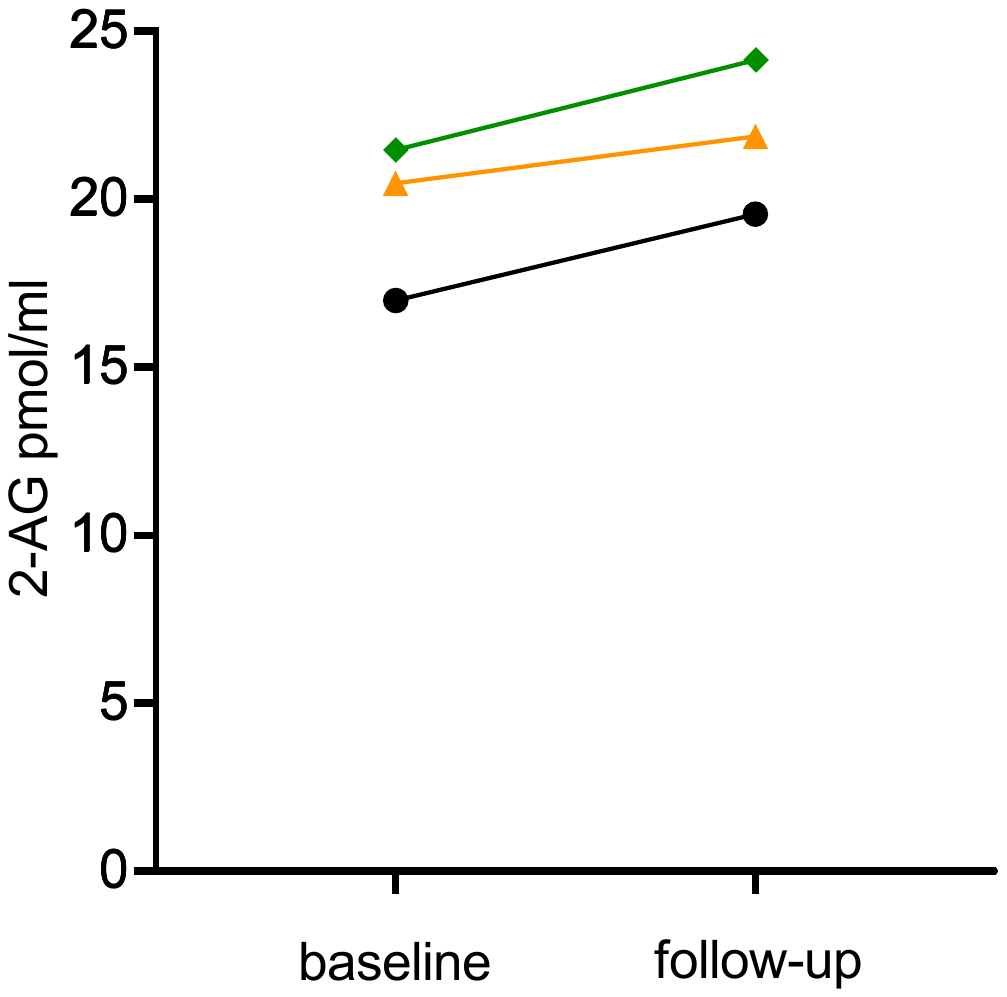

**Figure S2.** Elevated 2-arachidonylglycerol (2-AG) was significantly associated with lower cocaine craving scores measured by the cocaine craving questionnaire (CCQ) within **a)** recreational CU in dark blue but not in recreational CU is shown in light blue. This association was primarily driven by **b)** alcohol dependence and its interaction with 2-AG in dependent CU indicating that the relationship between 2-AG and cocaine craving was only seen in individuals showing current cocaine and co-morbid alcohol dependence shown in orange. Scatter plots reflects the negative association between 2-AG and the CCQ sum score, including trendline and 95% CI.

**a)**

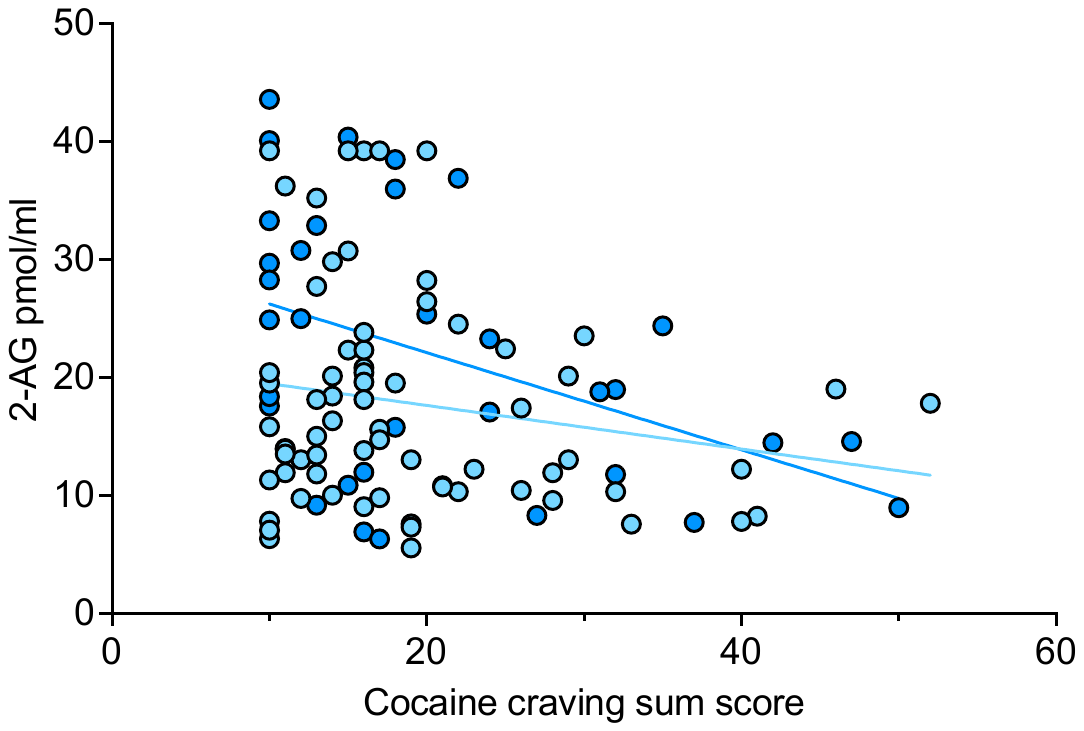

**b)**

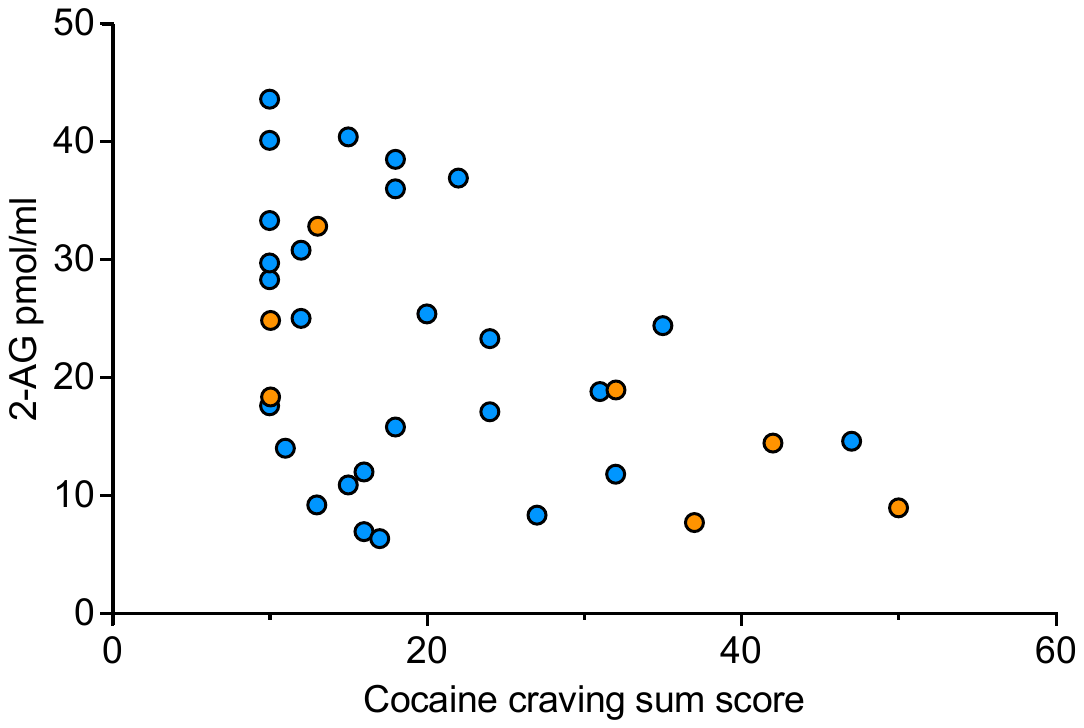

**Figure S3.** Within controls (n=16), basal plasma levels of 2-AG were not significantly associated with mGluR5 density of the **a)** amygdala **b)** thalamus **c)** hippocampal regions **d)** anterior cingulate cortex (ACC) **e)** midcingulate cortex (MCC) **f)** insula **g)** caudate.

a) b)

| Amygdala Vnorm | 2-AG pmol/ml |
| --- | --- |
| 1.17143 | 34.3 |
| 1.21362 | 28.5 |
| 1.36216 | 12.9 |
| 1.4178 | 31.8 |
| 1.21771 | 17.3 |
| 1.51097 | 37.7 |
| 1.42558 | 8.37 |
| 1.35403 | 16 |
| 1.25206 | 6.49 |
| 1.35096 | 9.13 |
| 1.61577 | 23 |
| 1.67261 | 5.99 |
| 2.05797 | 7.51 |
| 1.91378 | 15.6 |
| 1.38292 | 26.8 |
| 1.87425 | 11.6 |

| Thalamus Vnorm | 2-AG pmol/ml |
| --- | --- |
| 1.23626 | 34.3 |
| 1.20104 | 28.5 |
| 1.25966 | 12.9 |
| 1.27262 | 31.8 |
| 1.22001 | 17.3 |
| 1.33738 | 37.7 |
| 1.31474 | 8.37 |
| 1.37942 | 16 |
| 1.26184 | 6.49 |
| 1.24708 | 9.13 |
| 1.4499 | 23 |
| 1.43144 | 5.99 |
| 1.57624 | 7.51 |
| 1.75935 | 15.6 |
| 1.33846 | 26.8 |
| 1.52708 | 11.6 |

c) d)

| ACC Vnorm | 2-AG pmol/ml |
| --- | --- |
| 1.42602 | 34.3 |
| 1.38676 | 28.5 |
| 1.43443 | 12.9 |
| 1.57531 | 31.8 |
| 1.44902 | 17.3 |
| 1.72695 | 37.7 |
| 1.51214 | 8.37 |
| 1.78423 | 16 |
| 1.43967 | 6.49 |
| 1.44025 | 9.13 |
| 1.73273 | 23 |
| 1.87939 | 5.99 |
| 2.27115 | 7.51 |
| 2.2293 | 15.6 |
| 1.54344 | 26.8 |
| 2.12992 | 11.6 |

| Hippocampal region Vnorm | 2-AG pmol/ml |
| --- | --- |
| 1.2347 | 34.3 |
| 1.2249 | 28.5 |
| 1.28823 | 12.9 |
| 1.35217 | 31.8 |
| 1.32622 | 17.3 |
| 1.4127 | 37.7 |
| 1.36874 | 8.37 |
| 1.40502 | 16 |
| 1.25838 | 6.49 |
| 1.29967 | 9.13 |
| 1.57254 | 23 |
| 1.77788 | 5.99 |
| 1.96692 | 7.51 |
| 1.91828 | 15.6 |
| 1.36422 | 26.8 |
| 1.81777 | 11.6 |

e) f)

| MCC Vnorm | 2-AG pmol/ml |
| --- | --- |
| 1.37823 | 34.3 |
| 1.26429 | 28.5 |
| 1.32578 | 12.9 |
| 1.46896 | 31.8 |
| 1.35262 | 17.3 |
| 1.50138 | 37.7 |
| 1.31645 | 8.37 |
| 1.67422 | 16 |
| 1.35923 | 6.49 |
| 1.36838 | 9.13 |
| 1.57207 | 23 |
| 1.72255 | 5.99 |
| 1.87354 | 7.51 |
| 2.10566 | 15.6 |
| 1.43978 | 26.8 |
| 1.8625 | 11.6 |

| Insula Vnorm | 2-AG pmol/ml |
| --- | --- |
| 1.38864 | 34.3 |
| 1.36254 | 28.5 |
| 1.47302 | 12.9 |
| 1.5545 | 31.8 |
| 1.30371 | 17.3 |
| 1.67249 | 37.7 |
| 1.48568 | 8.37 |
| 1.61592 | 16 |
| 1.42348 | 6.49 |
| 1.44258 | 9.13 |
| 1.74338 | 23 |
| 1.88294 | 5.99 |
| 2.22762 | 7.51 |
| 2.19883 | 15.6 |
| 1.50789 | 26.8 |
| 1.98322 | 11.6 |

g)

| Caudate Vnorm | 2-AG pmol/ml |
| --- | --- |
| 1.37035 | 34.3 |
| 1.33256 | 28.5 |
| 1.42577 | 12.9 |
| 1.52704 | 31.8 |
| 1.32721 | 17.3 |
| 1.64815 | 37.7 |
| 1.5342 | 8.37 |
| 1.69768 | 16 |
| 1.49448 | 6.49 |
| 1.47278 | 9.13 |
| 1.87287 | 23 |
| 1.92753 | 5.99 |
| 2.21914 | 7.51 |
| 2.2296 | 15.6 |
| 1.55335 | 26.8 |
| 2.092 | 11.6 |
